# Salted roads lead to edema and reduced locomotor function in wood frogs

**DOI:** 10.1101/2021.03.23.436008

**Authors:** Lauren E. Frymus, Debora Goedert, Francisco Javier Zamora-Camacho, Peter C. Smith, Caroline J. Zeiss, Mar Comas, Timothy A. Abbott, Silvia P. Basu, Jason C. DeAndressi, Mia E. Forgione, Michael J. Maloney, Joseph L. Priester, Faruk Senturk, Richard V. Szeligowski, Alina S. Tucker, Mason Zhang, Ryan Calsbeek, Steven P. Brady

## Abstract

Human activities have caused massive losses of natural populations across the globe. Like many groups, amphibians have experienced substantial declines worldwide, driven by environmental changes such as habitat conversion, pollution, and disease emergence. Each of these drivers is often found in close association with the presence of roads. Here we report a novel consequence of roads affecting an amphibian native to much of North America, the wood frog (*Rana sylvatica*). Across 38 populations distributed from southern to central New England, we found that adult wood frogs living adjacent to roads had higher incidence and severity of edema (bloating caused by fluid accumulation) during the breeding season than frogs living away from the influence of roads. This effect was best explained by increased conductivity of breeding ponds, caused by runoff pollution from road salt used for de-icing. Edema severity was negatively correlated with locomotor performance in more northerly populations. Interestingly, northern populations experience more intense winters, which tends to result in more de-icing salt runoff and increased energetic demands associated with overwintering cryoprotection needs. Thus, this emerging consequence of roads appears to impose potential fitness costs associated with locomotion, and these effects might be most impactful on populations living in regions where de-icing is most intense.

## Introduction

Human activities are a leading driver of wild population declines, extirpations, and extinctions, culminating in the loss of global biodiversity (Ceballos et al. 2015, Leigh et al. 2019, Rosenberg et al. 2019). These losses shorten the branches of the tree of life and leave lasting if not permanent alterations to long-evolved interactions among species (Emer et al. 2019), modifying ecosystem function and the services they provide (Symstad et al. 1998, Cardinale et al. 2012).

Among the many groups of organisms declining across the globe, amphibians have captured a special place in the attention of researchers, fueling a surge in scientific inquiry over the past three decades (Green et al. 2020). Causes of declines in amphibian populations are complex and dependent on both regional context and species-specific adaptations, but unsurprisingly share a common theme of human activity (Grant et al. 2020). Drivers include habitat loss and alteration, pollution, climate change, and disease. These and many other human-mediated drivers can act synergistically, confronting populations with conditions far exceeding the capacity of existing trait variation or the potential for adaptive evolutionary or plastic responses (reviewed in Green et al. 2020).

Lurking among the numerous drivers of amphibian declines lies a common but often overlooked thread: roads. The planet is swathed in some 64 million km of roads (Central Intelligence Agency 2013), with 25 million km more expected by 2050 (Dulac 2013). This vast network contributes an extensive and generally negative set of impacts on natural populations (Trombulak and Frissell 2000, Fahrig and Rytwinski 2009), even as some may be adapting (Brady and Richardson 2017). Perhaps most direct and apparent, roads result in huge numbers of road kill – dispatching up to 1 million vertebrates per day in the U.S. alone (Lalo 1987) – a phenomenon that disproportionately affects amphibians (Glista et al. 2008). Roads can facilitate the spread of both disease (Urban 2006) and invasive species (Mortensen et al. 2009); they fragment habitats (Reed et al. 1996), reduce connectivity (Shepard et al. 2008), and affect dispersal and gene flow (Riley et al. 2006). Roads are also substantial sources of pollution. Leaching and stormwater runoff have deposited countless contaminants into the environment for decades (Huber et al. 2016). Many contaminants can reach toxic levels in soil, surface waters, and groundwater (Marsalek et al. 1999, Tang et al. 2013, McIntyre et al. 2018), and these effects can extend to habitats located well beyond the road (Forman and Deblinger 2000). Some contaminants are fleeting, broken down by chemical or biological processes, but others are long-lived or inert and tend to increase over time (Kaushal et al. 2005). In short, this singular and physically narrow change to the landscape caused by roads spurs numerous and wide-reaching drivers of population decline. Identifying and understanding these drivers and their effects is essential for conservation efforts.

In areas with cold winters, a chief pollutant threatening amphibian populations is road salt. Due to de-icing practices and the pervasiveness of roads, salt pollution of surface and groundwaters has become common and widespread (Mullaney et al. 2009, Corsi et al. 2010). Even in large waterbodies (e.g., permanent ponds, lakes), runoff pollution can result in substantial salt concentrations exceeding national water quality criteria for aquatic life (Dugan et al. 2017, Kaushal et al. 2018). In smaller waterbodies however (e.g., vernal pools, wetlands), levels can be many times higher, approaching brackish conditions typical of coastal waters, a stark contrast to the generally salt-free conditions of small, inland surface waters. For many pool-breeding amphibians, these smaller waters are critical sites for reproduction, where embryos and larvae develop before metamorphosing and dispersing into terrestrial habitats. Across the complex life-history cycle common to many amphibians, sensitivity to salt tends to be highest for embryos and larvae (Gordon and Tucker 1965, Uchiyama et al. 1990, Karraker et al. 2008, Albecker and McCoy 2017). Thus, pool-breeding amphibians tend to face salt pollution at peak sensitivity, and this timing often coincides with annual peaks in salt concentration that occur after winter. At high enough concentrations, salt causes outright mortality in amphibians (Sanzo and Hecnar 2006, Brady et al. 2017). At lower concentrations, sublethal effects are numerous. Aquatic-stage exposure to salt affects behavior (Denoël et al. 2010) and modifies growth and developmental rates (Dananay et al. 2015, Brady 2017), it increases malformations and susceptibility to disease (Karraker and Ruthig 2009, Brady 2013, Hall et al. 2020), and it causes ‘carry-over’ effects, negatively impacting post-metamorphic survival in the terrestrial environment (Dananay et al. 2015); all of which can contribute to observed population declines (e.g., Karraker et al. 2008).

In the present study, we investigated an additional and novel physiological consequence of roads: severe gross edema found in adult frogs breeding in polluted, roadside ponds. Although edema is recognized as a disorder in veterinary and disease literature (e.g., Hadfield and Whitaker 2005, Miller et al. 2011), its occurrence and causes in natural amphibian populations have been overlooked (but see Hall et al. 2017). For the wood frog (*Rana sylvatica*) – the focus of our study – temporary edema can be found among breeding adults in early spring, apparently resulting naturally from overwintering physiology. During spring thaw, the reversal of the process whereby water is sequestered out of cells to prevent intra-cellular ice formation, is thought to cause temporary edema (Kling et al. 1994, Irwin et al. 1999). However, our preliminary observations suggest that spring edema is more severe in populations from polluted, roadside ponds compared to those from unpolluted, woodland ponds.

Here, we quantified the severity of edema in wood frogs from 38 ponds in New England, where road salt pollution is common. We posited that road salt might be causing severe edema, and therefore asked whether variation in edema prevalence and severity correlates with road adjacency and water conductivity, and if edema prevalence and severity increases with age, a proxy for lifetime exposure. We also investigated directly the effect of roadside pond water on edema by estimating body mass gain for animals exposed to water from roadside versus woodland ponds and compared to spring water. Next, we used dissections and histological preparations to characterize the relation between edematous outward appearance and potential underlying signs and causes. Since edema could be associated with winter physiology and spring thaw, we also asked if blood glucose levels correlated with edema severity. Finally, we investigated a potential fitness cost of edema. Specifically, we asked how jumping performance, a commonly used fitness proxy for frogs, is affected by edema severity.

## Methods

### Natural history

Wood frogs are widespread in North America, ranging in distribution from the eastern southern Appalachian Mountains and extending north- and westward to the Alaska-Yukon region within the Arctic circle (Martof and Humphries 1959, Green et al. 2014). To survive winter conditions, particularly in higher latitudes, wood frogs rely on physiological adaptations to freezing (Storey and Storey 1988, Costanzo et al. 2015). Wood frogs typically use ephemeral or other small, fishless ponds for breeding in early spring. Across our study area, breeding occurs typically between early March and mid-April, lasting about 1-2 weeks in any given pond. Immediately prior to breeding, adults migrate from upland terrestrial habitat to mate in ephemeral ponds. Males amplex females and fertilize eggs externally during oviposition. Each female lays a single clutch as an egg mass containing approximately 800–1100 eggs. Embryos develop over 2–3 weeks before hatching and continue to develop as aquatic larvae throughout spring and early summer until they metamorphose into terrestrial juveniles, which disperse into upland habitat. Adults can live for 5-6 years (Berven 2009, Brady et al. 2019), and apart from annual breeding, tend to spend most of their lives in terrestrial habitat. Juvenile wood frogs show strong natal site fidelity and, as adults, strong breeding site fidelity (Berven and Grudzien 1990).

### Edema severity/prevalence and environmental variation

We quantified edema by compiling data for 935 individual frogs from 38 breeding ponds (putative populations, referred interchangeably as “populations” hereafter) in New England (N=20 roadside, 18 woodland; Fig. 1). Following Brady (2013), ponds were categorized as either ‘roadside’ (< 10-15 m from a paved road) or ‘woodland’ (> 150 m from any road). Frogs were captured during four breeding seasons in three different study areas (spanning from 41.2-43.8 degrees North), each containing a mix of roadside and woodland populations: 1) Southern Connecticut (N=9 roadside, 9 woodland ponds in 2018), 2) Northeastern Connecticut (N=6 roadside, 5 woodland ponds in 2010), and 3) the Upper Valley of Vermont and New Hampshire (N=5 roadside, 4 woodland ponds in 2016 and 2017). In Northeastern Connecticut, frogs were captured on their inbound breeding migration using drift fences with pitfall traps placed adjacent to breeding ponds. Frogs from these populations were therefore not exposed to pond water immediately prior to capture. In the other two study areas, drift fences with pitfalls were supplemented with minnow traps. Thus, frogs from these populations had mixed histories of exposure to pond water immediately before capture. Both types of traps were checked daily for new captures. Captured frogs were returned to the lab and dorsal photographs were taken within two days. Edema severity was scored from these photographs on a 1-3 integer scale, corresponding to 1) non-edematous, 2) moderate edema, and 3) severe edema. For the analysis of prevalence, an individual was considered edematous if it had an edema score of either 2 or 3. The following environmental variables were measured during the spring breeding / rearing seasons in a subset of ponds: specific conductivity (N=31 ponds), dissolved oxygen (N=29 ponds), and pH (N=18 ponds).

**Figure 1.**
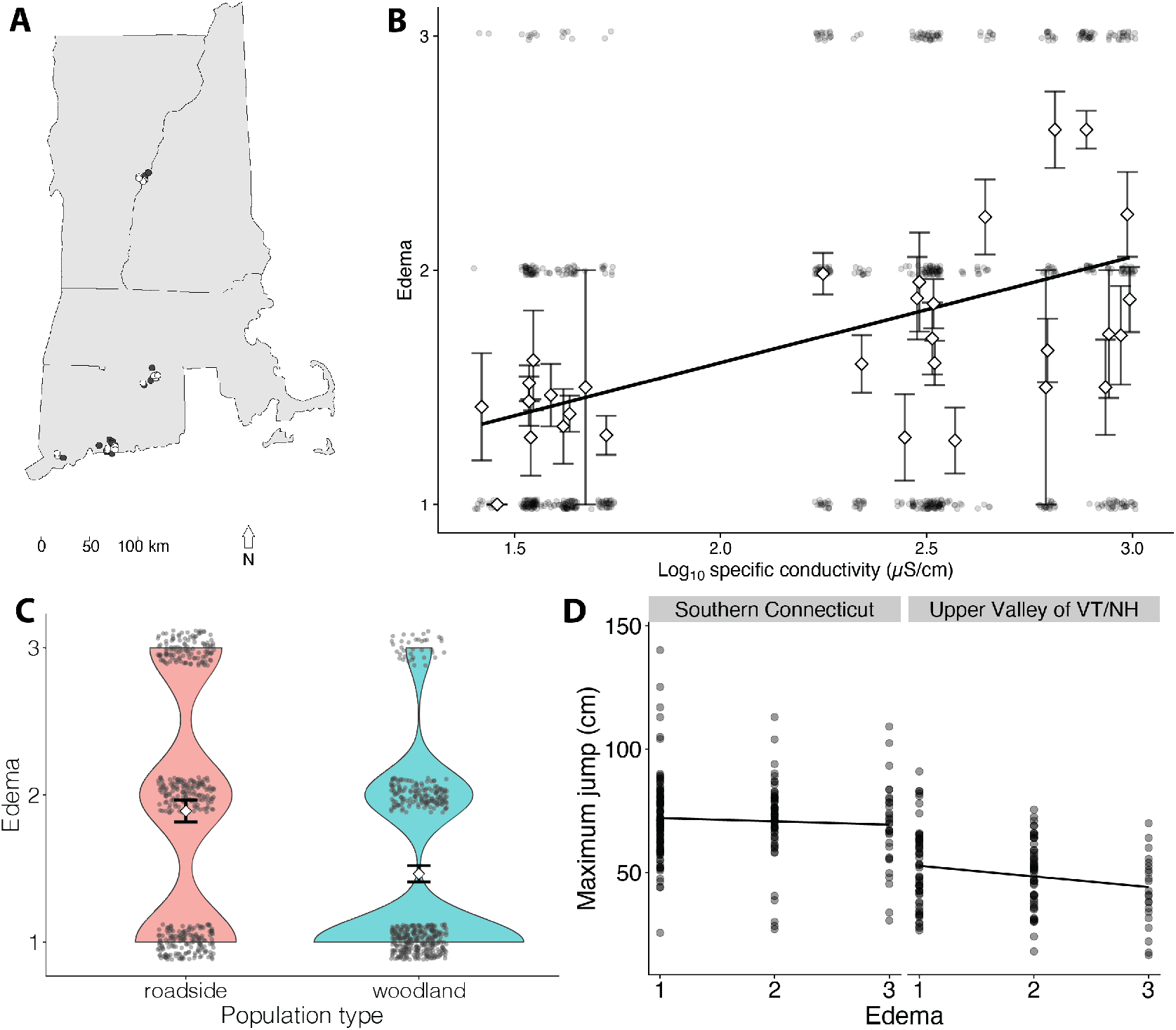
Study sites, edema prevalence, and jump performance. Panel A: Edema was scored in frogs from 38 ponds located within three regions in New England, from south to north: Southern Connecticut, Northeastern Conecticut, and the Upper Valley in Vermont/New Hampshire. Locations of roadside ponds (filled circles) and woodland ponds (open circles) are shown; points were jittered to reduce overlap. Panel B: Edema increased with increasing conductivity (a proxy for road salt). Diamonds (± 1 SE) represent pond-level average edema. Circles show individual data points, jittered to reduce overlap. Regression line was estimated with a linear model of pond-level averages for both edema and conductivity. Panel C: Edema was more severe in roadside ponds. Each point represents edema for a unique frog. Points are jiggered vertically and horizontally to reduce overlap. Violin plots show density distribution. Means (± 95% CI) for each population type are indicated by open diamonds. Panel D: Edema compromised jumping performance for (right) more northerly (*P* = 0.016) but not (left) more southerly populations (*P* = 0.479). Regression lines represent fixed effect of population type from mixed model.

We used separate mixed models to evaluate the effect of population type (roadside vs. woodland) on 1) edema severity score and 2) edema prevalence. Edema severity was fit using a linear model with a Gaussian distribution while edema prevalence was fit using a binomial model with a logit link. In each case, we first used AIC model selection to compare three candidate models fit with maximum likelihood and the following random effect structures: population, population and region (i.e. Southern Connecticut, Northeastern Connecticut, or Upper Valley of Vermont and New Hampshire), and population nested within region. Using a threshold for delta AIC value of 2, the model with the single random effect of population was preferred for both edema severity and prevalence. Fixed effect inference of severity was made after refitting the selected model with restricted maximum likelihood estimation. Here and elsewhere when linear mixed models were used, p-values were based on degrees of freedom calculated with Satterthwaite’s approach implemented in the R package ‘lmerTest’ (Kuznetsova et al. 2017). Because our edema severity data could potentially be considered ordinal, we also composed a mixed model using the R package ‘ordinal’ (Christensen 2019). Inference was similar between these two modeling approaches (*P* < 0.001 in each case) and we therefore interpreted results from the linear model. For the binomial responses of prevalence, inference was made using a likelihood ratio test between the selected model and a reduced model in which the response mean was used as the sole fixed effect in place of population type (i.e. intercept-only model). In a separate suite of standard linear models, we analyzed edema in response to conductivity, dissolved oxygen, and pH. Pond-level averages of edema score were used as the response variable in each model. Conductivity was log-transformed because of its wide range (26-984 µS/cm) and bimodal distribution between woodland and roadside ponds.

We analyzed the relation between adult age and both edema severity and prevalence using data from 74 individuals collected in 2014 from 9 populations (N=4 roadside, 5 woodland) in the Upper Valley of Vermont and New Hampshire. Individuals used in this analysis were collected during the inbound breeding migration (prior to arriving at ponds) using drift fences with pitfall traps. Edema was estimated from photos as described above and age was estimated using skeletochronology of toe clips (methods described in Brady et al. 2019). We used a linear model with either prevalence or severity as the response variable, and the interaction between age and population type for fixed effects.

### Exposure-induced change in mass

We examined the effects of pond water exposure on adult wood frog water uptake, a potential cause of edema. We analyzed data from a prior study conducted in the same Northeastern CT populations, designed originally to test the effects of early-embryo exposure to polluted pond water on larval survival (see Brady 2017). Briefly, adult frogs were captured during the breeding season using drift fences with pitfall traps adjacent to breeding ponds. All frogs were captured on their inbound migration and thus were not yet exposed to breeding pond water at the time of capture. At each breeding pond, captured frogs were measured for snout-vent length (SVL) and weighed prior to being paired haphazardly to breed in 5.1 L plastic aquaria filled with 1 L of either breeding-pond water or spring water. For roadside populations, this design resulted in approximately half the treatments being roadside water and the other half being spring water. For woodland populations, about half the treatments were woodland pond water and the other half were spring water. Across 11 ponds (N=6 roadside, 5 woodland), an average of 8.5 (range: 4-14) breeding pairs were stocked into each treatment (i.e. breeding-pond water or spring water). Exposure period was determined by the time required to breed. Following breeding, adults were re-weighed.

We analyzed change in body mass relative to SVL for males prior to and after breeding. Females were excluded from this analysis because change in mass caused by water is confounded with change in mass due to oviposition. Specifically, we calculated a body condition index (BCI) as mass per unit SVL (i.e. mass/SVL), both before and after exposure to pond or spring water that occurred while breeding. We then calculated delta BCI as the ratio of post-breeding BCI to pre-breeding BCI. Thus, delta BCI values > 1 indicate body mass gain whereas those < 1 indicate body mass loss. We used separate linear mixed models with population as the random effect to analyze both delta BCI and exposure duration across the interaction of population type and exposure water type. We used the R package ‘emmeans’ (Lenth et al. 2018) to estimate marginal means for each of these models and to compute 95% confidence intervals with respect to a null value of 1 (i.e. no change in body mass). We applied Tukey contrasts to make pairwise comparisons of both delta BCI and exposure duration across this interaction.

### Morphology, histology and blood glucose

In spring 2020, we necropsied six frogs from Southern Connecticut populations (N=1 with severe edema, N=5 non-edematous, all male) to identify the underlying source of edema and to evaluate potential changes in gross morphology associated with edema. Following prosection and gross dissection, soft tissues (including gastrointestinal tract, liver, spleen, kidneys, lung, heart, reproductive tissues and remainder of body) were fixed in 10% neutral buffered formalin, while long bones, associated muscle, nerve and skin, as well as brain and eyes (in skull), were fixed in Bouin’s solution for one week. Tissues were processed using standard paraffin processing sectioned at 5 µm, and stained with hematoxylin and eosin for light microscopic evaluation. Images were taken with a Zeiss Axioskop and Zeiss Axiocam MrC camera. We also measured blood glucose in subset of 54 male frogs from 6 roadside ponds (N = 35 frogs) and 4 woodland ponds (N = 19 frogs) in Southern Connecticut. We obtained a blood sample using a 30.5 gauge needle to pierce the facial vein (Forzán et al. 2012). A drop of blood was then placed on a testing strip and assayed using a Bayer Contour® Next blood glucose meter. We used a mixed model with population as a random effect to analyze blood sugar in response to edema and population type. We included body mass as a covariate in this analysis because of its potential effect on blood glucose irrespective of edema related to pollution. The two highest blood glucose observations were removed from analysis because they were 12 and 6.8 times the median absolute deviation.

### Jumping performance

To evaluate whether jumping performance might be compromised by edema, we conducted jump trials on a subset of frogs scored for edema from Southern Connecticut (N = 270 in 2019) and the Upper Valley of Vermont and New Hampshire (N = 258 in 2016 and 2017). Frogs were placed on one end of a 0.91 × 5 m piece of brown paper and at least 5 consecutive jumps were recorded, or frogs were placed between two plastic walls (height: 34 cm; length: 3.4 m) separated by approximately 30 cm, such that frogs could only jump forward. Frogs were motivated to jump as needed by tapping their urostyle with a finger (Brady et al. 2019). Marks were placed where frogs landed and the maximum distance between marks was recorded. We used a mixed model with population as the random effect to analyze the relation between jumping performance and edema. For each frog, we used maximum jump distance from among all jumps in a given trial as the response variable. We analyzed jump performance collectively and separately for the two regions. We also used a mixed model with population as the random effect to analyze whether jump performance varied between the two regions.

## Results

### Edema severity/prevalence and environmental variation

Across 38 ponds, edema was more severe and more prevalent in roadside than woodland ponds (Fig. 1). Specifically, model estimates of edema score were 1.84 in roadside ponds compared to 1.46 in woodland ponds (*F*_*1,30*.*0*_ = 14.77, *P* < 0.001). Variance between ponds (regardless of type) estimated from the random effect accounted for 12.3% of total edema severity variance. The probability of finding edema in roadside ponds was estimated at 0.605 compared to 0.367 in woodland ponds. Edema was therefore 65% more prevalent in roadside versus woodland ponds. Among environmental variables, edema severity was correlated with log-transformed conductivity (*F*_*1,29*_ *=* 17.72, *P* < 0.001; Fig 1) but not pH (*F*_*1,16*_ *=* 0.603, *P* = 0.45) or dissolved oxygen (*F*_*1,27*_ *=* 0.00, *P* = 0.996). Conductivity averaged 545 µS/cm in roadside ponds (range: 170-984 µS/cm), representing a 14-fold increase over the average 38 µS/cm (range: 26-53) found in woodland ponds (*F*_*1,29*_ *=* 38.57, *P* < 0.001).

Neither edema severity (*X*^*2*^ = 0.146, df = 1, *P* = 0.702) nor prevalence (*X*^*2*^ = 0.01, df = 1, *P* = 0.921) varied across the interaction of age and population type. Similarly, when main effects were considered, we found no effect of age or population type on edema severity (age: *F*_*1,71*_ *=* 1.85, *P* = 0.178; population type: *F*_*1,71*_ *=* 0.207, *P* = 0.651) or prevalence (age: *X*^*2*^ = 0.957, df = 1, *P* = 0.328; population type: *X*^*2*^ = 0.003, df = 1, *P* = 0.958).

### Exposure-induced change in mass

Delta BCI, or the change in body mass relative to SVL, varied across the interaction of water type X population type (Fig. 2; *F*_*1,339*.*3*_ *=* 18.02, *P* < 0.001). Specifically, wood frogs exposed to roadside water had the highest relative change in mass compared to those in all other treatments. Model estimates indicated that roadside wood frogs exposed to roadside water maintained their mass, whereas wood frogs in all other combinations of water and population type lost mass (Supplemental Table 1). The mean duration of exposure was 5.94 days (range of 1-13 days) and differed across the interaction of water type X population type (*F*_*1,290*.*23*_ *=* 3.97, *P* = 0.047). Tukey post-hoc pairwise comparisons revealed that the only difference in exposure time was between woodland populations in woodland water versus spring water (*P* = 0.026; woodland populations exposure duration: 5.91 days for woodland pond water compared to 4.96 days for spring water; all other pairwise comparisons: *P* > 0.505), indicating differences in exposure time did not drive the effect found in delta BCI for roadside populations.

**Figure 2.**
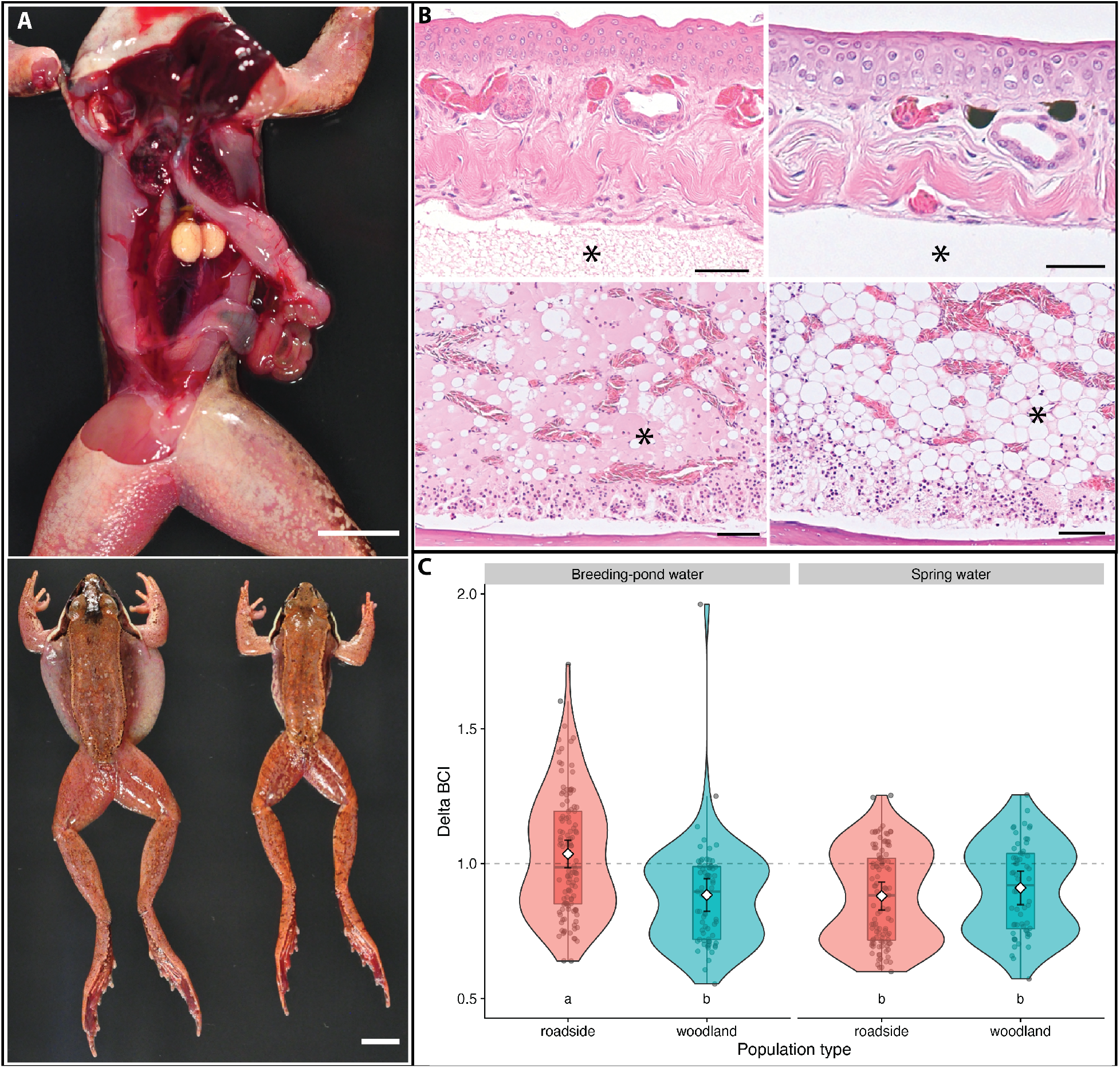
Edema and water balance. (A) On gross dissection of the edematous frog (top), edema fluid rapidly dissipated from the subcutis (not visible), and was evident to a much lesser degree in the coelomic cavity. Testes were well developed. Marked subcutaneous edema is evident in an edematous frog (bottom left) compared to a non-edematous frog (bottom right). Bars = 1cm. (B) On histology, the grossly edematous animal exhibited subcutaneous edema (asterisk, top left) not evident in the non-edematous animal (asterisk, top right). Marked serous atrophy of fat was evident in bone marrow of the edematous animal (asterisk, bottom left) in comparison to non-edematous animals with greater retention of intact adipocytes within bone marrow (asterisk, bottom right). Bars = 50µm, hematoxylin and eosin. (C) Change in adult male mass relative to body size (i.e. body condition index, or ‘BCI’) during a controlled breeding experiment is shown for different exposure waters. Mass was measured before and after breeding. Delta BCI of 1 (dashed line) indicates no change. Experimental exposure to (far left) roadside pond water caused frogs from roadside ponds gain or maintain water on average during the breeding period. In contrast, on average, frogs from woodland ponds exposed to (left center) woodland water lost mass. Similarly, individuals exposed to spring water – whether from (right center) roadside ponds or (far right) woodland ponds – lost mass on average. Raw data are shown as gray circles. Violin plots and boxplots are overlain. White diamonds and error bars represent estimated marginal means and corresponding 95% confidence intervals.

### Morphology, histology and blood glucose

On gross examination, marked subcutaneous edema was evident in the frog with severe edema (Fig. 2). Upon dissection, edema fluid rapidly dissipated from the subcutis, and was evident to a much lesser degree in the coelomic cavity. In all frogs, the gastrointestinal tract was devoid of ingesta or digesta, no fat stores were present, and testes were well developed. On histology, the grossly edematous animal exhibited subcutaneous edema. Adipose tissue was absent, and bone marrow was characterized by marked serous fat atrophy (or ‘gelatinous bone marrow transformation’) a condition associated with starvation and characterized by loss of both fat and red-blood stem (‘hematopoietic’) cells, and the deposition of extracellular mucous (Fig. 2). In contrast, grossly non-edematous animals had absent or mild subcutaneous edema and greater retention of intact adipocytes within bone marrow. Remaining tissues were unremarkable, with the exception of a trematode segment characterized by an acoelomic body, tegument and intestinal tract within the liver on a non-edematous animal. This was considered an incidental finding. In all animals, spermatogenesis was vigorous.

Blood glucose varied with respect to edema severity (*F*_*1,43*.*8*_ *=* 7.00, *P* = 0.011) and body mass (*F*_*1,27*.*4*_ *=* 5.06, *P* = 0.033) but not population type (*F*_*1,10*.*0*_ *=* 0.05, *P* = 0.831). When accounting for population type and mass, along with random effects of population, blood glucose increased 13.16 mg/dL with each one-unit increase in edema score and decreased 4.2 mg/dL for each gram of body mass.

### Jumping performance

When all populations from the two regions with jumping data were analyzed together, edema was not correlated with maximum jumping distance in (male) frogs (*F*_*1,183*.*2*_ *=* 0.278, *P* = 0.097). However, separate analyses for each region revealed that edema negatively affected maximum jump distance for frogs in the Upper Valley (*F*_*1,131*_ *=* 4.92, *P* = 0.016) but not for frogs in Southern Connecticut (*F*_*1,199*.*4*_ *=* 0.50, *P* = 0.479). Specifically, maximum jump distance for Upper Valley frogs declined by 4.3 cm for each unit increase in edema. Thus, on their best jumps, the frogs with severe edema jumped approximately 9 cm less than non-edematous frogs. Jumping performance also varied between regions independent of edema (*F*_*1,11*.*7*_ *=* 71.05, *P* < 0.001). Southern Connecticut frogs jumped an estimated 71 cm compared to 49 cm for Upper Valley frogs. Thus, maximum jump for Upper Valley frogs was about 68% that of Southern Connecticut frogs.

## Discussion

Edema was both more prevalent and more severe in roadside populations subject to salt runoff compared to unsalted woodland populations across three different regions in the northeastern USA. In support of salt runoff being a causative agent of edema, we found that edema severity was correlated with conductivity in breeding ponds, and that frogs experimentally exposed to roadside pond water during breeding increased fluid retention. Edema also correlated with increased blood glucose, as well as with decreased jumping performance in northerly (but not southerly) populations. Together, these results reveal that pollution from roads appears to be increasing the severity of edema in amphibians, and that this relatively unexplored consequence of roads imposes a potential fitness cost through its effect on locomotion.

Our gross morphological and histological findings indicate that fluid accumulation in the subcutis was indeed responsible for the edematous appearance of frogs. Several factors identified from the veterinary literature are known to cause subcutaneous edema, including fungal, bacterial, or viral infection, exposure to low solute water, and kidney diseases (reviewed by Pessier 2009). Interestingly, high salinity water might in some instances help prevent some diseases in amphibians (e.g., Stockwell et al. 2015). However, the consequences of salt runoff for disease susceptibility in amphibians are complex and not fully understood, with some studies suggesting that osmotic stress caused by salt might be detrimental to the survival of individuals in populations with ranavirus infections (Hall et al. 2020). Although we cannot completely rule out ranavirus as a factor, infected individuals typically present ulcerative and hemorrhagic syndromes in multiple tissues (Cunningham et al. 1996, Docherty et al. 2003), which we did not observe in the edematous animal dissected or among the animals scored for edema.

In our study, exposure to low solute water can definitively be ruled out as a possible factor, since animals presenting edema were more frequently those exposed to high solute water. Frogs have highly permeable skin and must osmoregulate in aquatic environments. For frogs in freshwater, the tendency is for water to move from the environment into the frog. Curiously, from an osmotic perspective, adding salt to freshwater should decrease the osmotic potential of the external environment relative to that of the frog. That is, by adding ions to the pond, water pressure exerted on frog skin decreases. In contrast, in unpolluted woodland ponds, external water pressure is higher than that found in polluted roadside ponds. Thus, based purely on osmotic pressure, there should be a greater tendency for woodland, not roadside, water to move into frogs. And yet we found edema was more severe in salty, roadside environments, pointing to dysfunction in osmoregulation. Therefore, it is likely that the culprit of edema is kidney disorder, potentially incurred from osmoregulatory stress from prior exposure to high salinity.

Increased edema severity in roadside ponds might also be linked to overwintering physiology. Beginning in fall as temperatures decline, wood frogs accumulate urea (Costanzo and Lee 2005) and continue to increase liver glycogen stores. During freezing, glycogen is metabolized into glucose (Costanzo and Lee 2013, Costanzo et al. 2015) and distributed to cells and organs. Water is moved to the coelomic cavity and lymph system (Lee et al. 1992), and cytosol moves osmotically to extracellular spaces. These physiological changes help prevent cells from freezing while permitting coelomic and subdural ice formation to occur with little damage to organs and tissue. Estimates indicate that up to about 2/3 of body water can become frozen in wood frogs during the winter (Layne Jr and Lee Jr 1987, Storey and Storey 1988, Costanzo and Lee 2013). Thawing in the spring can be rapid and appears to be a source of fleeting edema (Kling et al. 1994, Irwin et al. 1999) during which time there might be a lag period for water resorption from coelomic and subdural spaces, consistent with our histological and gross dissection findings. Although it is unclear why road proximity might increase severity of thaw-induced edema, conceivably, pollution might trigger a hyper-reactive freeze tolerance response and/or a reduced capacity for resorption. In which case, it is not surprising that we found elevated levels of glucose in the spring, suggesting either that edematous frogs have an amplified cryoprotective response (concentrating more glucose in their cells) and/or they metabolize glucose more slowly after thawing.

Regardless of whether edema is linked to winter physiology or a direct consequence of kidney disease, the severity we detected in roadside populations prompts consideration of what conditions in roadside environments increase edema severity. For instance, is edema exacerbated by the aquatic environment, terrestrial environment, or both? For adults in our study, one such exposure could have occurred during embryonic and larval stages (as wood frogs show strong natal site fidelity; Berven and Grudzien 1990) while another exposure could have occurred in adult stages but in prior breeding years (as wood frogs also show strong breeding site fidelity; Berven and Grudzien 1990). In either case, some degree of carry-over effect is implied. Exposure could also occur in terrestrial roadside habitats, for instance through consumption of saltier food items or exposure to saltier soils.

The edema we describe occurred in breeding frogs captured either on route to or soon after arriving in their breeding ponds. This means that for at least a subset of frogs, the onset of edema occurred before they were exposed to pond water. This finding indicates that edema sets in either during the short migration from upland hibernacula to breeding ponds, or that frogs are edematous prior to migration. In late fall, frogs tend to choose overwintering sites close to their breeding ponds (O’Connor and Rittenhouse 2016), such that salt runoff exposure in the terrestrial environment might be exacerbating edema that forms naturally following the reversal of winter cryoprotection. Interestingly, we found no relation between age and edema severity or prevalence, suggesting that a single exposure to the roadside environment might be just as likely to generate edema as a history of adult breeding in roadside ponds.

Our study additionally suggests a significant impact of salt runoff exposure in the aquatic environment. We found higher edema severity with increased conductivity of the aquatic environment, a variable widely used to measure road salt pollution. In New England, like many temperate regions with cold winters, roadside ponds have become salinized due to runoff from road salt (Kaushal et al. 2005, Karraker et al. 2008, Brady 2012, Dugan et al. 2017). Although a variety of salts are used in de-icing practices, NaCl is the agent applied most abundantly, both in our study region and throughout the U.S. (Mullaney et al. 2009). Conductivity values in roadside ponds averaged 545 µS/cm, and individual values ranged up to 2050 µS/cm. Based on prior calibrations (G. Benoit unpublished data), we estimate these values correspond to 300 and 1128 mg/L of total NaCl, or 182 mg/L and 684 mg/L of Cl-, respectively. Moreover, we found that frogs experimentally exposed to roadside water during breeding maintained mass on average while those exposed to woodland or spring water lost mass. We interpret this change in mass as being driven primarily by water gain or loss, thus representing a relative increase in water retention or uptake in roadside water, because frogs did not have access to food during this period. Saltier aquatic environments should result in frogs taking up more ions for osmoregulation (Greenwald 1972), which could lead to higher water retention (Park and Do 2020) consistent with our findings (Fig. 2). For example, Park and Do (2020) found that exposure of black-spotted frogs (*Pelophylax nigromaculatus*) to about 9,400 µS/cm resulted in increased blood electrolytes and that frogs became edematous. The authors attribute this effect to renal dysfunction resulting from the increase in electrolytes. Future studies would be required to disentangle the relative importance of pre- and post-metamorphosis exposure, as well as exposure through aquatic versus terrestrial environments.

Finally, we found that jumping performance was compromised by edema in the more northerly populations of Vermont and New Hampshire, but not those of Southern Connecticut. Here again, overwintering conditions might mediate this effect. For instance, in northern populations, more severe and longer winters should produce more intense cryoprotective physiological changes (Costanzo et al. 2015). This investment in cryoprotection might bear a cost to muscle performance needed for locomotion, consistent with our findings that Upper Valley frogs jumped just 68% the distance of Southern Connecticut frogs. Costanzo et al. (2015) report that muscle catabolism is an important component of freeze tolerance observed in northern (Alaska) wood frog populations but absent in Ohio populations, and that it results in skeletal muscle atrophy. The authors suggest that this mechanism is essential to increase nitrogen availability for urea production given the more extreme winters in Alaska. Conceivably, harsher winters in Vermont and New Hampshire induce this same mechanism and cause muscle atrophy that diminishes jumping performance. Also, the serous fat atrophy we found with severe edema suggests that edematous frogs consume more energy for cryoprotection. Given the importance of energy and locomotion during spring breeding, particularly in the context of road crossings and swimming performance during mating, these consequences could bear impacts on fitness components including survival and reproductive success and, therefore, on the viability of higher latitude populations.

In this study, we investigated possible causes and fitness consequences of increased edema severity, representing an amphibian population threat from roads and runoff pollution that has remained largely unexplored. Increased edema severity in wood frogs appears to be a wide-ranging phenomenon in road-adjacent habitats, driven by road salt pollution. Given the regional distribution of edema reported here, we expect that this phenomenon likely affects wood frogs across much of their range, although the consequences of edema might be most severe for populations experiencing more intense winters. Interestingly, northern areas where winters are more intense tend to use more road salt. Thus, the presence of these multiple, correlated stressors suggests that wood frogs in more northerly populations might be both most likely to experience higher edema severity near roads and locomotor consequences as a result. Many important questions remain about the mechanisms and consequences of severe edema in roadside wood frogs. Indeed, while road salt is a prevalent contaminant and almost certainly the most common pollutant by mass in these ponds, other contaminants could be contributing to edema, such as heavy metals and aromatic hydrocarbons, and such effects might be interactive. Identifying the causes of severe edema and its impacts on populations should guide future investigations.

## Acknowledgements

This work was supported by Mianus River Gorge Preserve, Elm City Innovation Collaborative, Yale Institute for Biospheric Studies, EEES Graduate fellowship and Cramer funds, Guarini School of Graduate and Advanced Studies McCulloch Fellowship, CAPES graduate fellowship (SwB 13442/13-9), and the National Science Foundation (DEB #1011335, DEB #1655092). We thank D. Skelly for project advice, and J. Peterman and S. Rodrigues for help in the field and lab. We are grateful to the many private land owners and public land managers who allowed us to work on their lands.

## Literature Cited

Albecker, M. A., and M. W. McCoy. 2017. Adaptive responses to salinity stress across multiple life stages in anuran amphibians. Frontiers in zoology 14:40–40.

Berven, K. A. 2009. Density dependence in the terrestrial stage of wood frogs: evidence from a 21-year population study. Copeia 2009:328–338.

Brady, S. P. 2012. Road to evolution? Local adaptation to road adjacency in an amphibian (Ambystoma maculatum). Scientific Reports 2.

Brady, S. P. 2013. Microgeographic maladaptive performance and deme depression in response to roads and runoff. PeerJ 1:e163.

Brady, S. P. 2017. Environmental exposure does not explain putative maladaptation in road-adjacent populations. Oecologia 184:931–942.

Brady, S. P., and J. L. Richardson. 2017. Road ecology: shifting gears toward evolutionary perspectives. Frontiers in Ecology and the Environment 15:91–98.

Brady, S. P., J. L. Richardson, and B. K. Kunz. 2017. Incorporating evolutionary insights to improve ecotoxicology for freshwater species. Evolutionary Applications 10:829–838.

Brady, S. P., F. J. Zamora-Camacho, F. A. Eriksson, D. Goedert, M. Comas, and R. Calsbeek. 2019. Fitter frogs from polluted ponds: The complex impacts of human-altered environments. Evolutionary Applications 12:1360–1370.

Cardinale, B. J., J. E. Duffy, A. Gonzalez, D. U. Hooper, C. Perrings, P. Venail, A. Narwani, G. M. Mace, D. Tilman, and D. A. Wardle. 2012. Biodiversity loss and its impact on humanity. Nature 486:59–67.

Ceballos, G., P. R. Ehrlich, A. D. Barnosky, A. García, R. M. Pringle, and T. M. Palmer. 2015. Accelerated modern human–induced species losses: Entering the sixth mass extinction. Science Advances 1:e1400253.

Central Intelligence Agency. 2013. The World Factbook 2012-13. Central Intelligence Agency.

Christensen, R. H. B. 2019. Regression Models for Ordinal Data [R package ordinal version 2019.12-10].

Corsi, S., D. Graczyk, S. Geis, N. Booth, and K. Richards. 2010. A fresh look at road salt: aquatic toxicity and water-quality impacts on local, regional, and national scales. Environmental Science & Technology 44:7376–7382.

Costanzo, J. P., and R. E. Lee. 2005. Cryoprotection by urea in a terrestrially hibernating frog. Journal of Experimental Biology 208:4079–4089.

Costanzo, J. P., and R. E. Lee. 2013. Avoidance and tolerance of freezing in ectothermic vertebrates. Journal of Experimental Biology 216:1961–1967.

Costanzo, J. P., A. M. Reynolds, M. C. F. do Amaral, A. J. Rosendale, and R. E. Lee Jr. 2015. Cryoprotectants and extreme freeze tolerance in a subarctic population of the wood frog. PLoS ONE 10:e0117234.

Cunningham, A., T. S. Langton, P. Bennett, J. Lewin, S. Drury, R. Gough, and S. Macgregor. 1996. Pathological and microbiological findings from incidents of unusual mortality of the common frog (Rana temporaria). Philosophical Transactions of the Royal Society of London. Series B: Biological Sciences 351:1539–1557.

Dananay, K. L., K. L. Krynak, T. J. Krynak, and M. F. Benard. 2015. Legacy of road salt: Apparent positive larval effects counteracted by negative postmetamorphic effects in wood frogs. Environmental Toxicology and Chemistry 34:2417–2424.

Denoël, M., M. Bichot, G. F. Ficetola, J. Delcourt, M. Ylieff, P. Kestemont, and P. Poncin. 2010. Cumulative effects of road de-icing salt on amphibian behavior. Aquatic Toxicology 99:275–280.

Docherty, D. E., C. U. Meteyer, J. Wang, J. Mao, S. T. Case, and V. G. Chinchar. 2003. Diagnostic and molecular evaluation of three iridovirus-associated salamander mortality events. Journal of Wildlife Diseases 39:556–566.

Dugan, H. A., S. L. Bartlett, S. M. Burke, J. P. Doubek, F. E. Krivak-Tetley, N. K. Skaff, J. C. Summers, K. J. Farrell, I. M. McCullough, A. M. Morales-Williams, D. C. Roberts, Z. Ouyang, F. Scordo, P. C. Hanson, and K. C. Weathers. 2017. Salting our freshwater lakes. Proceedings of the National Academy of Sciences:201620211.

Dulac, J. 2013. Global land transport infrastructure requirements. Paris: International Energy Agency 20:2014.

Emer, C., M. Galetti, M. A. Pizo, P. Jordano, and M. Verdú. 2019. Defaunation precipitates the extinction of evolutionarily distinct interactions in the Anthropocene. Science Advances 5:eaav6699.

Fahrig, L., and T. Rytwinski. 2009. Effects of roads on animal abundance: an empirical review and synthesis. Ecology and Society 14.

Forman, R. T. T., and R. D. Deblinger. 2000. The ecological road-effect zone of a Massachusetts (USA) suburban highway. Conservation Biology 14.

Forzán, M. J., R. V. Vanderstichel, C. T. Ogbuah, J. R. Barta, and T. G. Smith. 2012. Blood collection from the facial (maxillary)/musculo-cutaneous vein in true frogs (family Ranidae). J Wildl Dis 48:176–180.

Glista, D. J., T. L. DeVault, and J. A. DeWoody. 2008. Vertebrate road mortality predominantly impacts amphibians. Herpetological conservation and Biology 3:77–87.

Gordon, M. S., and V. A. Tucker. 1965. Osmotic regulation in the tadpoles of the crab-eating frog (Rana cancrivora). Journal of Experimental Biology 42:437–445.

Grant, E. H. C., D. A. Miller, and E. Muths. 2020. A Synthesis of Evidence of Drivers of Amphibian Declines. Herpetologica.

Green, D. M., M. J. Lannoo, D. Lesbarrères, and E. Muths. 2020. Amphibian population declines: 30 years of progress in confronting a complex problem. Herpetologica 76:97–100.

Green, D. M., L. A. Weir, G. S. Casper, and M. J. Lannoo. 2014. North American amphibians: distribution and diversity. Univ of California Press.

Greenwald, L. 1972. Sodium Balance in Amphibians from Different Habitats. Physiological Zoology 45:229–237.

Hadfield, C. A., and B. R. Whitaker. 2005. Amphibian emergency medicine and care. Pages 79-89 in Seminars in Avian and Exotic Pet Medicine. Elsevier.

Hall, E. M., S. P. Brady, N. M. Mattheus, R. L. Earley, M. Diamond, and E. J. Crespi. 2017. Physiological consequences of exposure to salinized roadside ponds on wood frog larvae and adults. Biological Conservation 209:98–106.

Hall, E. M., J. L. Brunner, B. Hutzenbiler, and E. J. Crespi. 2020. Salinity stress increases the severity of ranavirus epidemics in amphibian populations. Proceedings of the Royal Society B: Biological Sciences 287:20200062.

Huber, M., A. Welker, and B. Helmreich. 2016. Critical review of heavy metal pollution of traffic area runoff: Occurrence, influencing factors, and partitioning. Science of the Total Environment 541:895–919.

Irwin, J. T., J. P. Costanzo, and J. Lee, Richard E. 1999. Terrestrial hibernation in the northern cricket frog, Acris crepitans. Canadian Journal of Zoology 77:1240–1246.

Karraker, N. E., J. P. Gibbs, and J. R. Vonesh. 2008. Impacts of road deicing salt on the demography of vernal pool-breeding amphibians. Ecological Applications 18:724–734.

Karraker, N. E., and G. R. Ruthig. 2009. Effect of road deicing salt on the susceptibility of amphibian embryos to infection by water molds. Environmental Research 109.

Kaushal, S. S., P. M. Groffman, G. E. Likens, K. T. Belt, W. P. Stack, V. R. Kelly, L. E. Band, and G. T. Fisher. 2005. Increased salinization of fresh water in the northeastern United States. Proceedings of the National Academy of Sciences of the United States of America 102:13517–13520.

Kaushal, S. S., G. E. Likens, M. L. Pace, R. M. Utz, S. Haq, J. Gorman, and M. Grese. 2018. Freshwater salinization syndrome on a continental scale. Proceedings of the National Academy of Sciences 115:E574–E583.

Kling, K., J. Costanzo, and R. Lee. 1994. Post-freeze recovery of peripheral nerve function in the freeze-tolerant wood frog, Rana sylvatica. Journal of Comparative Physiology B 164:316–320.

Kuznetsova, A., P. B. Brockhoff, and R. H. Christensen. 2017. lmerTest package: tests in linear mixed effects models. Journal of statistical software 82:1–26.

Lalo, J. 1987. The problem of road kill. American forests (USA).

Layne Jr, J. R., and R. E. Lee Jr. 1987. Freeze tolerance and the dynamics of ice formation in wood frogs (Rana sylvatica) from southern Ohio. Canadian Journal of Zoology 65:2062–2065.

Lee, R. E., J. P. Costanzo, E. C. Davidson, and J. R. Layne Jr. 1992. Dynamics of body water during freezing and thawing in a freeze-tolerant frog (Rana sylvatica). Journal of Thermal Biology 17:263–266.

Leigh, D. M., A. P. Hendry, E. Vázquez-Domínguez, and V. L. Friesen. 2019. Estimated six per cent loss of genetic variation in wild populations since the industrial revolution. Evolutionary Applications 12:1505–1512.

Lenth, R., H. Singmann, J. Love, P. Buerkner, and M. Herve. 2018. Emmeans: Estimated marginal means, aka least-squares means. R package version 1:3.

Marsalek, J., Q. Rochfort, B. Brownlee, T. Mayer, and M. Servos. 1999. An exploratory study of urban runoff toxicity. Water science and technology 39:33–39.

Martof, B. S., and R. L. Humphries. 1959. Geographic Variation in the Wood Frog Rana sylvatica. American Midland Naturalist 61:350–389.

McIntyre, J. K., J. I. Lundin, J. R. Cameron, M. I. Chow, J. W. Davis, J. P. Incardona, and N. L. Scholz. 2018. Interspecies variation in the susceptibility of adult Pacific salmon to toxic urban stormwater runoff. Environmental Pollution 238:196–203.

Miller, D., M. Gray, and A. Storfer. 2011. Ecopathology of ranaviruses infecting amphibians. Viruses 3:2351–2373.

Mortensen, D. A., E. S. Rauschert, A. N. Nord, and B. P. Jones. 2009. Forest roads facilitate the spread of invasive plants. Invasive Plant Science and Management 2:191–199.

Mullaney, J. R., D. L. Lorenz, and A. D. Arntson. 2009. Chloride in groundwater and surface water in areas underlain by the glacial aquifer system, northern United States. U.S. Geological Survey Scientific Investigations Report 2009–5086.

O’Connor, J. H., and T. A. Rittenhouse. 2016. Snow cover and late fall movement influence wood frog survival during an unusually cold winter. Oecologia 181:635–644.

Park, J. K., and Y. Do. 2020. Physiological response of Pelophylax nigromaculatus adults to salinity exposure. Animals (Basel) 10:1698.

Pessier, A. P. 2009. Edematous frogs, urinary tract disease, and disorders of fluid balance in amphibians. Journal of Exotic Pet Medicine 18:4–13.

Reed, R. A., J. Johnson-Barnard, and W. L. Baker. 1996. Contribution of roads to forest fragmentation in the Rocky Mountains. Conservation Biology 10:1098–1106.

Riley, S. P., J. P. Pollinger, R. M. Sauvajot, E. C. York, C. Bromley, T. K. Fuller, and R. K. Wayne. 2006. FAST-TRACK: A southern California freeway is a physical and social barrier to gene flow in carnivores. Molecular Ecology 15:1733–1741.

Rosenberg, K. V., A. M. Dokter, P. J. Blancher, J. R. Sauer, A. C. Smith, P. A. Smith, J. C. Stanton, A. Panjabi, L. Helft, M. Parr, and P. P. Marra. 2019. Decline of the North American avifauna. Science 366:120.

Sanzo, D., and S. J. Hecnar. 2006. Effects of road de-icing salt (NaCl) on larval wood frogs (Rana sylvatica). Environmental Pollution 140.

Shepard, D. B., A. R. Kuhns, M. J. Dreslik, and C. A. Phillips. 2008. Roads as barriers to animal movement in fragmented landscapes. Animal Conservation 11:288–296.

Stockwell, M., J. Clulow, and M. Mahony. 2015. Evidence of a salt refuge: chytrid infection loads are suppressed in hosts exposed to salt. Oecologia 177:901–910.

Storey, K. B., and J. M. Storey. 1988. Freeze tolerance in animals. Physiological Reviews 68:27–84.

Symstad, A. J., D. Tilman, J. Willson, and J. M. Knops. 1998. Species loss and ecosystem functioning: effects of species identity and community composition. Oikos:389–397.

Tang, J. Y., R. Aryal, A. Deletic, W. Gernjak, E. Glenn, D. McCarthy, and B. I. Escher. 2013. Toxicity characterization of urban stormwater with bioanalytical tools. Water Research 47:5594–5606.

Trombulak, S. C., and C. A. Frissell. 2000. Review of ecological effects of roads on terrestrial and aquatic communities. Conservation Biology 14:18–30.

Uchiyama, M., T. Murakami, and H. Yoshizawa. 1990. Notes on the development of the crab-eating frog, Rana cancrivora. Zoological Science 7:p73–78.

Urban, M. C. 2006. Road facilitation of trematode infections in snails of northern Alaska. Conservation Biology 20:1143–1149.

