## Supplemental Table 1 for "Salted roads lead to edema and reduced locomotor function in wood frogs"

**Supplemental Table 1.** Results of mixed model showing estimates of exposure-induced change in mass (i.e. ‘delta BCI’).

*Population type X water type	Estimate	df	Lower-upper 95% CL*

roadside X roadside water	1.036	14.5	0.985-1.087

woodland X woodland water	0.884	22.4	0.823-0.944

roadside X spring water	0.880	15.7	0.828-0.932

woodland X spring water	0.910	25.8	0.848-0.972
